# Sucrose-responsive osmoregulation of plant cell size by a long non-coding RNA

**DOI:** 10.1101/2024.02.19.581113

**Authors:** Jakub Hajný, Tereza Trávníčková, R. M. Imtiaz Karim Rony, Sebastian Sacharowski, Michal Krzyszton, David Zalabák, Christian S. Hardtke, Aleš Pečinka, Szymon Swiezewski, Jaimie M. van Norman, Ondřej Novák

**Author notes:** Corresponding author: Jakub Hajný.

## Abstract

The shoot of green plants is the primary site of carbon assimilation into sugars, the key source of energy and metabolic building blocks. The systemic transport of sugars is essential for plant growth and morphogenesis. Plants evolved intricate networks of molecular players to effectively orchestrate the subcellular partitioning of sugars. Dynamic distribution of these osmotically active compounds is a handy tool to regulate cell turgor pressure. Pressure-induced mechanical forces play an instructive role in developmental biology across kingdoms. Here, we functionally characterized a long non-coding RNA, *CARMA,* as a negative regulator of a receptor-like kinase, CANAR. Sugar-responsive *CARMA* specifically fine-tunes *CANAR* expression in the phloem, the route of sugar transport. By controlling sugar distribution, the CARMA-CANAR module allows cells to flexibly adapt to the external osmolality and adjust the size of vascular cell types during organ growth and development. We identify a nexus of plant vascular tissue formation with cell internal pressure monitoring and reveal a novel functional aspect of long non-coding RNAs in developmental biology.

## Introduction

In contrast to the circulatory vascular system of vertebrates, plants evolved non-circulatory specialized vascular bundles with two distinct long-distance transport routes. The xylem is a unidirectional root-to-shoot path for the transport of water and minerals from the soil. The phloem route transports carbon assimilates, amino acids, RNAs, and hormones from source tissues (e.g. mature leaves) into sink tissues (such as juvenile leaves, roots, meristems, and reproductive organs) (*1*, *2*). The hydrostatic pressure differences between source and sink drive the flow of the phloem content (*3*). In most plants, sucrose is the main form of assimilated carbon from photosynthesis, making it the central metabolite in plant growth and development. Sucrose is synthesized from fructose and glucose in photosynthetically active cells. Plants favor non-reducing sugar sucrose since high concentrations of reducing sugars can non-enzymatically glycosylate essential proteins and interfere with their functionality (*4*). Sucrose export from photosynthetic cells (mesophyll in leaves) to the apoplast is facilitated by SUGARS WILL EVENTUALLY BE EXPORTED TRANSPORTERS (SWEETs) efflux proteins. Then, sucrose enters the phloem via SUCROSE TRANSPORTERs (SUCs), a process termed apoplastic phloem loading. SUCs are H^+^/sucrose symporters, loading sucrose against its concentration gradient. In sink tissues, sucrose is unloaded from the phloem and distributed via SWEET proteins. Sink tissues convert sucrose back to glucose and fructose in the apoplast by cell wall-bound invertase enzymes. Ultimately, the sugars are consumed or stored in vacuoles (*4*, *5*).

Plant growth involves physical remodeling of cell wall mechanics and cell hydrostatic pressure. Plant cells have a high intracellular hydrostatic pressure, called turgor pressure, which results from water uptake in response to the solute concentration (e.g. ions and sugars) and is counterbalanced by the rigid yet dynamic cell walls (*6*, *7*). If osmotic conditions change, plant cells regulate water and ion transport across the plasma membrane (PM) and remodel their cell wall to compensate for the turgor pressure difference. The balance between turgor pressure and cell wall tension at the cell level translates to the tissue level, driving tissue patterning. These mechanical forces play an instructive role in developmental biology across kingdoms. For example, accumulating evidence suggests that in the shoot, the epidermis possesses thicker cell walls, providing a high resistance pillar for aerial organ development. In the root, the endodermis likely plays a similar role as the epidermis in the shoot. Both internal turgor pressure and external mechanical perturbations can alter cell size, geometry, polarity, cell division plane orientations, and, thus, the final plant shape (*8*).

In the *Arabidopsis thaliana* root, INFLORESCENCE AND ROOT APICES RECEPTOR KINASE (IRK), a leucine-rich repeat receptor-like kinase (LRR-RLK) regulates stele (i.e., the vascular cylinder surrounded by the pericycle layer) size, and restricts excessive endodermal cell divisions (*9*). IRK’s closest homolog PXY/TDR-CORRELATED 2 (PXC2), also called CANALIZATION-RELATED RECEPTOR-LIKE KINASE (CANAR), exerts an overlapping, partially redundant function despite not being expressed in the same tissues (*10*). Recently, IRK and CANAR/PXC2 were also reported to contribute to vascular patterning via auxin canalization (*10*, *11*). Interestingly, the relative number of cells in the stele between wild type (WT) and *CANAR* mutant/overexpression lines are similar despite the significant change in root stele area (*10*). This suggests mechanical remodeling, which, ultimately, alters cell volume instead of cell number. How CANAR participates in cell volume adjustment remains unknown.

Long non-coding RNAs (lncRNAs) are essential regulatory elements of eukaryotic transcriptomes. lncRNAs are versatile regulators of gene expression, functioning at different cellular levels often providing adaptive mechanisms to various stimuli (*12*). To date, only a handful of lncRNAs have been functionally characterized and implicated in aspects of plant development (*13*). In this study, we characterized a newly annotated lncRNA, *CARMA* (*<u>CA</u>NA<u>R</u> <u>M</u>ODUL<u>A</u>TOR IN PROTOPHLOEM*), which is located in the proximal promoter region of *CANAR* in *Arabidopsis thaliana*. *CARMA* fine-tunes the phloem-specific expression of *CANAR* in response to sucrose availability. Appropriate *CANAR* levels in the phloem are required to control cell size in the stele, suggesting the CARMA-CANAR module adjusts cell turgor in response to the environment to optimize cell size.

## Results

### Newly annotated antisense long non-coding RNA is located in the CANAR proximal promoter

We set out to unravel the molecular mechanisms regulating CANAR activity by re-examining its expression pattern. Previously, the transcriptional fusion of the entire intergenic region (4.7 kbp) upstream of the *CANAR* start codon with an ER-targeted green fluorescent protein (*pCANAR::erGFP)* showed weak activity in the Arabidopsis root tip (*10*). To observe a more native expression pattern, we rebuilt the reporter, adding the 3’ untranslated region (UTR) downstream of the *CANAR* stop codon to nuclear-targeted GFP and β-glucuronidase (*pCANAR::NLS-GFP-GUS-ter*). This reporter exhibited a markedly stronger fluorescent signal, localized mainly to the lateral root cap (LRC) and xylem (X), corresponding with the previous report (*10*). Lower expression could also be seen in the root phloem precursors: developing protophloem sieve elements (PPh) and metaphloem (MPh) (Fig. 1A). β-glucuronidase staining recapitulated previous observations (*14*), showing expression throughout the seedling vasculature. Staining in the first leaves occurred at the position of the future vasculature strands (Fig. 1B), supporting the previously described role of CANAR in vascular patterning via auxin canalization (*11*).

**Fig. 1.**
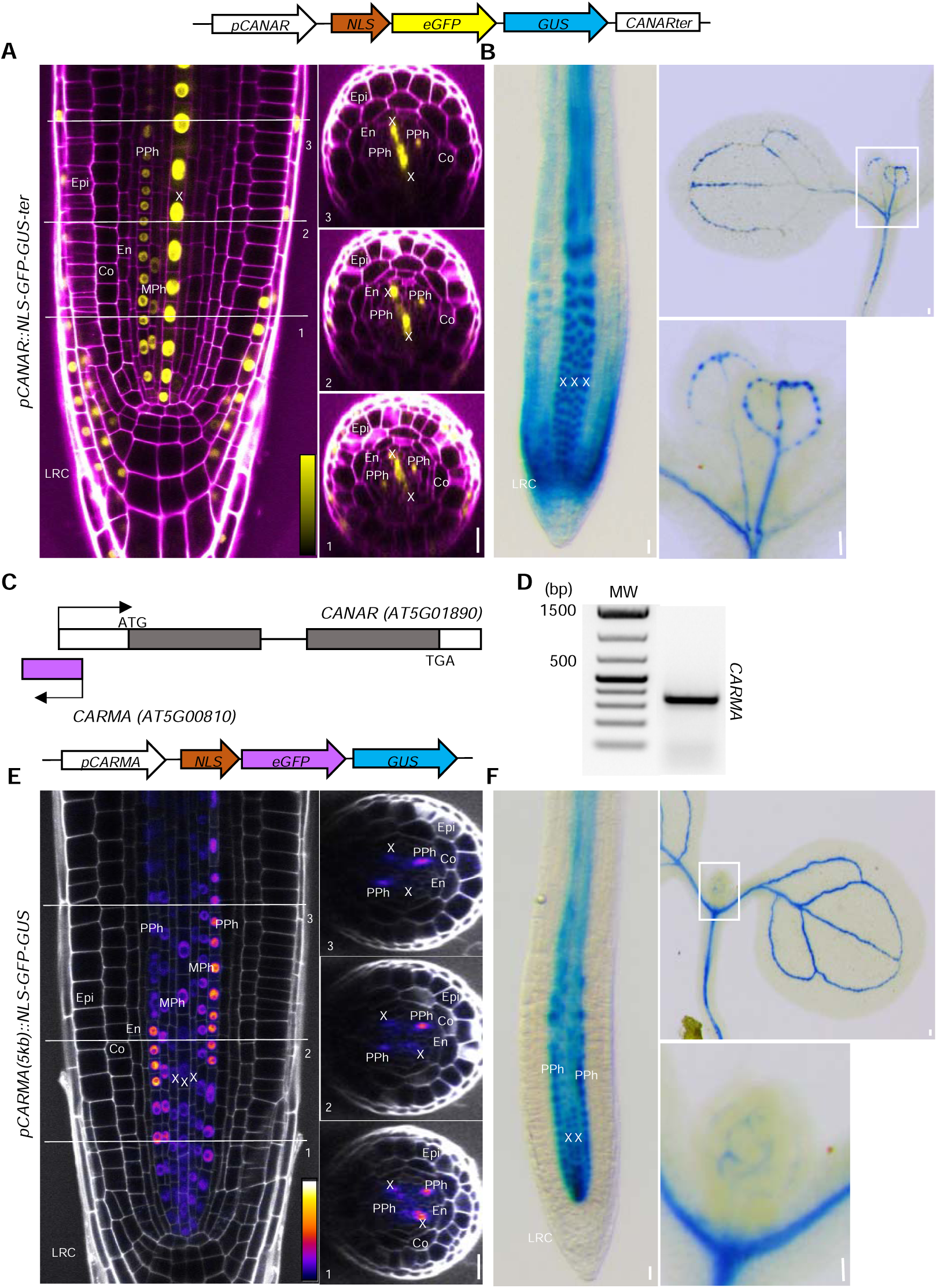
*CARMA* has a complementary expression with *CANAR* in root protophloem. (A) Confocal images of a primary root stained with propidium iodide (magenta) expressing *pCANAR::NLS-GFP-GUS-ter* (schematic depicted above images), shows *pCANAR* activity in xylem (X), developing protophloem sieve elements (PPh), lateral root cap (LRC), and with weaker expression in metaphloem precursors (MPh). (B) *pCANAR* activity in roots (left) and cotyledons and true leaves (right, leaves enlarged) visualized by β-glucuronidase (GUS) staining (blue). (C) A graphical representation of the *CARMA-CANAR* genomic locus. (D) sqRT-PCR of *CARMA* RNA from 5-day-old seedlings. (E) Confocal images of a primary root stained with propidium iodide (grey) expressing *pCARMA(5kb)::NLS-GFP-GUS* (depicted above images), showing *pCARMA* activity predominantly in PPh with weaker activity in MPh and X. (F) *pCARMA* activity in roots (left) and cotyledons and true leaves visualized by β-glucuronidase (GUS) staining (blue). Numbers in medial longitudinal confocal images represent the position of the transverse optical section taken from a Z-stack. For each reporter, ≥10 roots were examined. Scale bars 20 μm. Other cell types: Epi-epidermis, Co-cortex, En-endodermis.

During the design of *pCANAR* reporter, we noticed a newly annotated 353 bp antisense long non-coding RNA (lncRNA) (AT5G00810) in the proximal promoter region of *CANAR*, partially overlapping with its 5’ UTR (Fig. 1C). We hypothesized that this lncRNA, named *CARMA* (*<u>CA</u>NA<u>R</u> <u>M</u>ODUL<u>A</u>TOR IN PROTOPHLOEM*), might help us understand the relationship between tissue-specific expression of *CANAR* and its developmental functions. Using a semi-quantitative Reverse Transcription Polymerase Chain Reaction (sqRT-PCR), we confirmed that *CARMA* is expressed in seedlings and that the transcript is presumably polyadenylated as it could be amplified from oligo dT primed cDNA (Fig. 1D). We performed 5’ and 3’ Rapid Amplification of cDNA Ends (RACE) to define the full-length *CARMA* transcript. The transcription start site (TSS) largely matched annotation, whereas the 3’ end has several transcription termination sites (TTS). The annotated length of 353 bp constituted ∼50% of all *CARMA* transcripts with a maximum detected transcript length of 491 bp (Fig. S1A, B).

A transcriptional reporter containing 5 kb upstream of *CARMA* fused with *NLS-GFP-GUS* (*pCARMA(5kb)::NLS-GFP-GUS*), revealed *pCARMA* activity in the PPh with occasional expression in MPh. Additionally, in the meristematic zone, a shootward gradient of weaker expression in the xylem was also observed (Fig. 1E, F, S1C). The activity of *pCARMA* in the xylem was not seen with a shortened version of the promoter (*pCARMA(1.3kb)::NLS-GFP-GUS*) (Fig. S1D, E). Similar to *pCANAR, pCARMA* activity in the first leaves occurred at the position of the future vasculature strands, a manifestation of auxin canalization (*15*) (Fig. 1F). Thus, *pCANAR* and *pCARMA* have overlapping patterns of activity, but their intensity profiles are inverse suggesting a possible role for *CARMA* in transcriptional regulation of *CANAR*.

### CARMA controls leaf vascular patterning

*CARMA* expression in the cotyledons and first leaves prompted us to test the involvement of *CARMA* in leaf vascular patterning, a proxy for auxin canalization (*15*). We isolated an available T-DNA insertion loss-of-function mutant (*carma-1*) (Fig. S2A-C). Because the *carma-1* T-DNA insertion is close to the *CANAR* 5’ UTR (Fig. S2A), we tested whether it affects *CANAR* transcription. *CANAR* mRNA levels were unaffected (Fig. S2C), excluding the possibility of T-DNA-mediated knock-down of *CANAR*. Next, we generated transgenic lines overexpressing *CARMA* under the control of the constitutive cauliflower mosaic virus *35S* promoter (Fig. S2D). Two independent *35S::CARMA* overexpression lines showed a higher incidence of extra vascular loops, extra branches, and disconnections in the upper loops as compared to the wildtype (Col-0) control (Fig. 2A, B). These higher complexity venation phenotypes resembled that of *canar* mutants (*11*). In contrast, *carma-1* plants exhibited simpler venation, indicated by missing loops (Fig. 2C, D), similar to *35S::CANAR-GFP* (*11*).

**Fig. 2.**
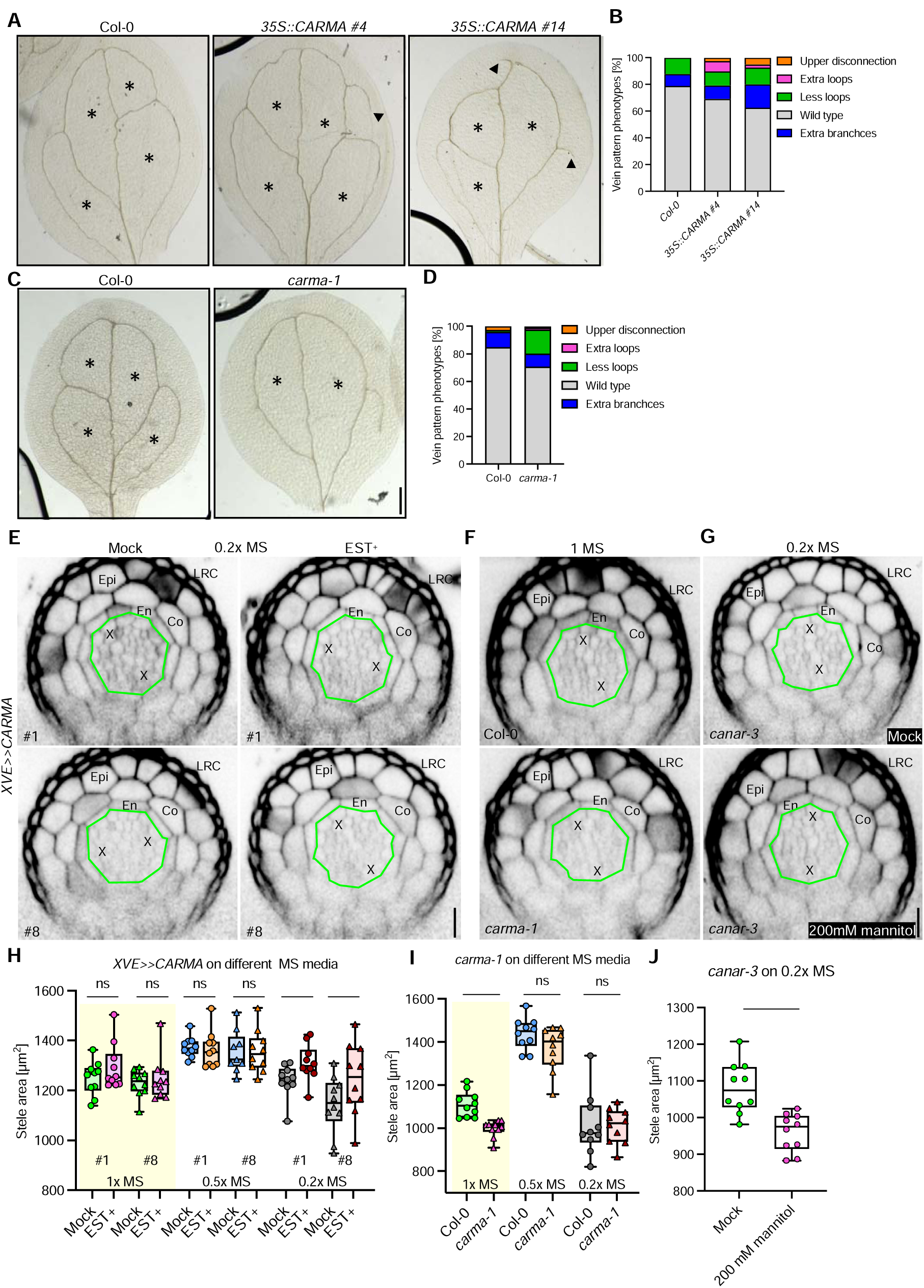
*CARMA* regulates leaf vascular patterning and root stele area. (A) and (C) representative images of cotyledon vasculature from 10-day-old Col-0, two independent *35S::CARMA* transgenic lines, and *carma-1* seedlings. Scale bars, 5 mm. (B) and (D) quantification of observed vein pattern phenotype as a percent. Black asterisks mark a number of closed loops. Black arrowheads highlight extra branches. For each genotype, ≥39 cotyledons were analyzed. (E) Transverse optical sections of 5-day-old root meristems stained with propidium iodide (gray) from two independent inducible CARMA overexpression (*XVE>>CARMA*) lines on 0.2x MS medium with (EST+) and without (mock) β-estradiol. (F) and (G) Transverseal optical sections of 5-day-old root meristems stained with propidium iodide of Col-0 and *carma-1* mutant on 1x MS and of *canar-3* on 0.2x MS medium supplemented with 200 mM mannitol. The outer edge of the stele area is indicated in green. (H) Box plots showing stele area quantification of *XVE>>CARMA* on different concentrations of MS mediums. (I) and (J) Box plots showing stele area quantifications of (F) and (G), respectively. Whiskers indicate max/min, box shows the interquartile range with a black line showing the median. Colored symbols are measurements from individual roots. The experiments were carried out three times (8-10 roots for each genotype per experiment); one representative biological replicate is shown. A one-way ANOVA test compared marked datasets (*P<0.05 and ****P<0.0001). Scale bars, 20 μm. Cell types: Epi-epidermis, Co-cortex, En-endodermis, X-xylem, LRC-lateral root cap. The transverse optical sections were taken approximately 100 µm from QC (quiescent center).

Together, the inverse intensity of *pCANAR* and *pCARMA* activity in the X/PPh and the opposite vein patterning phenotypes indicate that *CARMA* is a negative regulator of *CANAR* activity.

### CARMA mediates cell size changes in response to media osmolality in the stele

Whereas *canar-3* roots had an enlarged stele area, *CANAR* overexpression had the opposite effect. The stele area variance was due to a change in cell size and not cell number. This phenotype was conditional, manifested only in more hypotonic growth conditions where the agar plates contained 0.2x strength Murashige and Skoog medium (MS) basal salts media (*10*), suggesting involvement of internal water pressure in *CANAR* phenotype. Thus, we tested whether *CARMA* also plays a role in stele area control on media with different osmolality (0.2x, 0.5x, and 1x MS). As *35S* promoter activity is weak in the root meristem vasculature, we overexpressed *CARMA* under the β-estradiol inducible promoter (*16*) (*XVE>>CARMA)* (Fig. S2E). After β-estradiol treatment from germination onward, we observed a significantly enlarged stele area on 0.2x MS in two independent *XVE>>CARMA* lines compared to the Mock (Fig. 2E, H). Similar to what has been observed for *canar* mutant (*10*). Conversely, the *carma-1* roots exhibited a smaller stele area than WT, but only on 1x MS media (Fig. 2F, I), analogous to but weaker than *XVE>>CANAR* overexpression stele phenotype (*10*). Again, no change in the vascular cell number was observed (Fig. S2G, H; S3A, B), indicating the difference in stele area can be attributed to altered cell size, not proliferation. By measuring the distance from the endodermis to the lateral root cap, we confirmed that cell expansion is specific to the stele (Fig. S2F, S3C). Also, no change in root meristem length was observed (Fig. S2I, J; S3D, E), indicating that the stele area phenotype is not the result of changes in differentiation.

The conditional nature of these stele area phenotypes indicates a dependence on the osmolality of the media. Because the *canar-3* mutant has an enlarged stele on hypotonic media, we hypothesized that stele cells may retain excess water, making them bulkier. If true, lowering the intracellular water content would revert the phenotype. Indeed, the *canar-3* mutant grown on 0.2x MS media containing 200 mM of the osmolyte mannitol had decreased stele area compared to the Mock (Fig. 2G, J).

### CARMA fine-tunes CANAR expression in the root protophloem

The antisense orientation of *CARMA*, its inverse intensity expression profile in the X/PPh, and opposite leaf vasculature and stele area phenotypes with respect to *CANAR* imply that *CARMA* is a negative regulator of *CANAR*. To understand how *CARMA* influences CANAR function, we generated a set of transcriptional reporters consisting of the full-length 4.7 kbp *CANAR* promoter-*pCANAR::NLS-GFP-GUS-ter*, a partial deletion of *CARMA-pCANAR_CARMA*Δ*::NLS-GFP-GUS-ter,* and complete deletion of *CARMA* (removing part of the *CANAR* 5’ UTR as well)-*pCANAR_CARMA*ΔΔ*::NLS-GFP-GUS-ter* (Fig. 3A), transformed into *carma-1* plants. Using confocal microscopy, we observed that both deletions resulted in a significant, tissue-specific increase of *pCANAR* activity in the PPh to a level comparable to X. The insertional character of these transgenic lines does not allow absolute quantification; therefore, we opted for relative quantification of the PPh/X ratio of the fluorescence signal. Two independent transgenic lines were analyzed for each reporter (Fig. 3A, B; Fig. S4A, B). The similar outcomes of the *CARMA*Δ and *CARMA*ΔΔ deletions confirmed that changes in *pCANAR* activity are not due to an indirect impact of its partial 5’ UTR deletion.

**Fig. 3.**
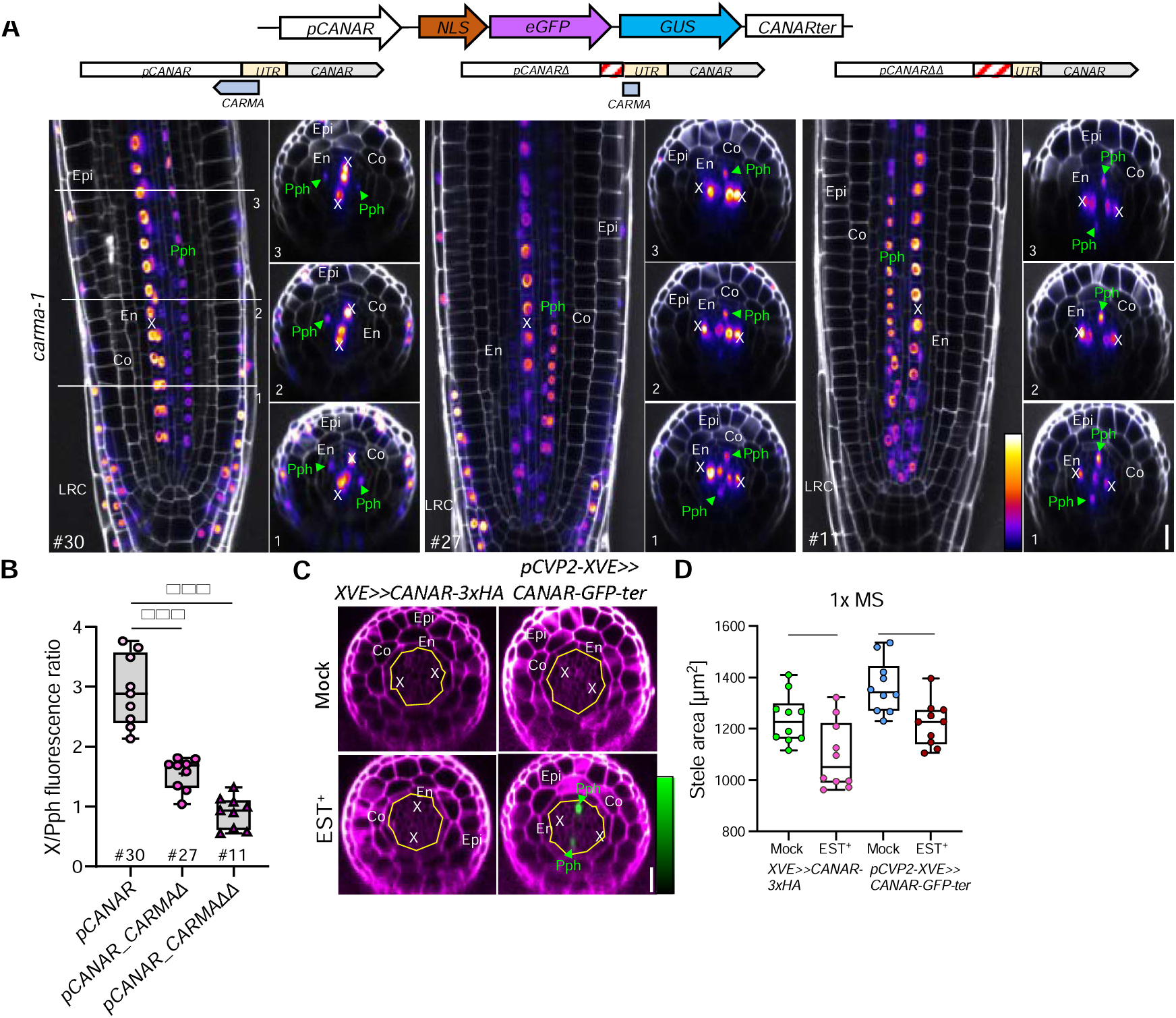
*CARMA* fine-tunes protophloem-specific expression of *CANAR*. (A) Representative confocal images of primary roots stained with propidium iodide (white) of *carma-1* plants expressing *pCANAR::NLS-GFP-GUS-ter, pCANAR_CARMA*Δ*::NLS-GFP-GUS-ter,* or *pCANAR_CARMA*ΔΔ*::NLS-GFP-GUS-ter* reporters (schematics shown above images). Both partial and complete deletion of *CARMA* led to increased *pCANAR* activity in the Pph (highlighted with a green label and arrowhead). Numbers represent the position of a transverse optical section taken from Z-stacks. These images were acquired using comparable settings. (B) Box plot showing relative fluorescence of reporters in (A) where the signal in the PPh is normalized to that in the X (see the Material and Method section for details). Whiskers indicate the max/min, the box shows the interquartile range, and the median is shown with a black line. Colored symbols show measurements for individual roots. (C) Transverse optical sections of 5-day-old root meristems stained with propidium iodide (magenta) from plants expressing *XVE>>CANARx3HA* and *pCVP2>>CANAR-GFP-ter* grown on 1x MS medium with (EST+) and without (mock)r β-estradiol from the time of germination. The outer edge of the stele is indicated by the yellow line. (D) Box plot showing stele area quantification of the plants in (C). Whiskers indicate the max/min, the box shows an interquartile range, and the median is shown with a black line. Colored symbols are the measurements from individual roots. These experiments were done three times (8-10 roots for each genotype per experiment); one representative biological replicate is shown. A one-way ANOVA test compared marked datasets (*P<0.05, **P<0.01, and ***P<0.001). Scale bars, 20 μm. Cell types: Epi-epidermis, Co-cortex, En-endodermis, PPh-developing protophloem sieve elements, X-xylem, and LRC-lateral root cap.

Our results demonstrate that *CARMA* modulates *CANAR* levels to establish high *CANAR* expression in X and relatively low in PPh. To address the biological significance of this stringent PPh-specific fine-tuning mechanism, we expressed *CANAR* either ubiquitously or tissue-specifically in the PPh. We utilized an *XVE>>CANAR-3xHA* line, which inducibly overexpresses *CANAR*, causing a marked decrease in the stele area (*10*). We could elicit this phenotype on 1x MS medium (Fig. 3C, D), where the *carma-1* plants exhibited a smaller stele area as well (Fig. 2F, I). Next, we generated *pCVP2>>XVE::CANAR-GFP-ter*, allowing for protophloem-specific inducible overexpression of *CANAR* (*17*). These transgenic plants grown on β-estradiol showed protophloem-specific GFP fluorescence (Fig. 3C) and had significantly decreased stele area, although not to the extent of *XVE>>CANAR-3xHA* (Fig. 3D). This could indicate that the xylem-expressed *CANAR* is also involved or it is a consequence of *CANAR* misexpression. Alternatively, the phenotypic difference might be due to the missing *CANAR* expression in MPh when the *CVP2* (*COTYLEDON VASCULAR PATTERN 2*) promoter is used.

Our results suggest that fine-tuned levels of *CANAR* in the PPh are required for the cell size adjustment in response to changes in external osmolality and are, thus, required for the optimization of stele area.

### CARMA mediates CANAR upregulation by sucrose

To better understand CANAR function, we set out to analyze the translational fusion of CANAR driven by its native promoter (*pCANAR::CANAR-GFP)* (*10*). Since the expression was too weak, we deployed a similar approach as with the *pCANAR::NLS-GFP-GUS-ter* transcriptional reporter, where the addition of the *CANAR* 3’UTR enhanced the fluorescence signal. Indeed, *pCANAR::CANAR-GFP-ter* provided a stronger signal (Fig. 4A). We noticed that fluorescence intensity and PM-localized signal in two independent transgenic lines depended on the presence of sucrose in the growth media. The effect was not observed after treatment with mannitol, a non-PM permeable sugar (Fig. 4A; S5A), or NaCl treatment, or changing the media osmolality (0.2x, 0.5x and 1x MS) (Fig. S5C). Three-fold higher sucrose concentration did not stimulate further increased accumulation of CANAR (Fig. S5B), indicating a maximum threshold. We further tested whether monosaccharides could exert a similar effect on CANAR expression. Indeed, the upregulation of CANAR was also observed after glucose treatment (Fig. S5B).

**Fig. 4.**
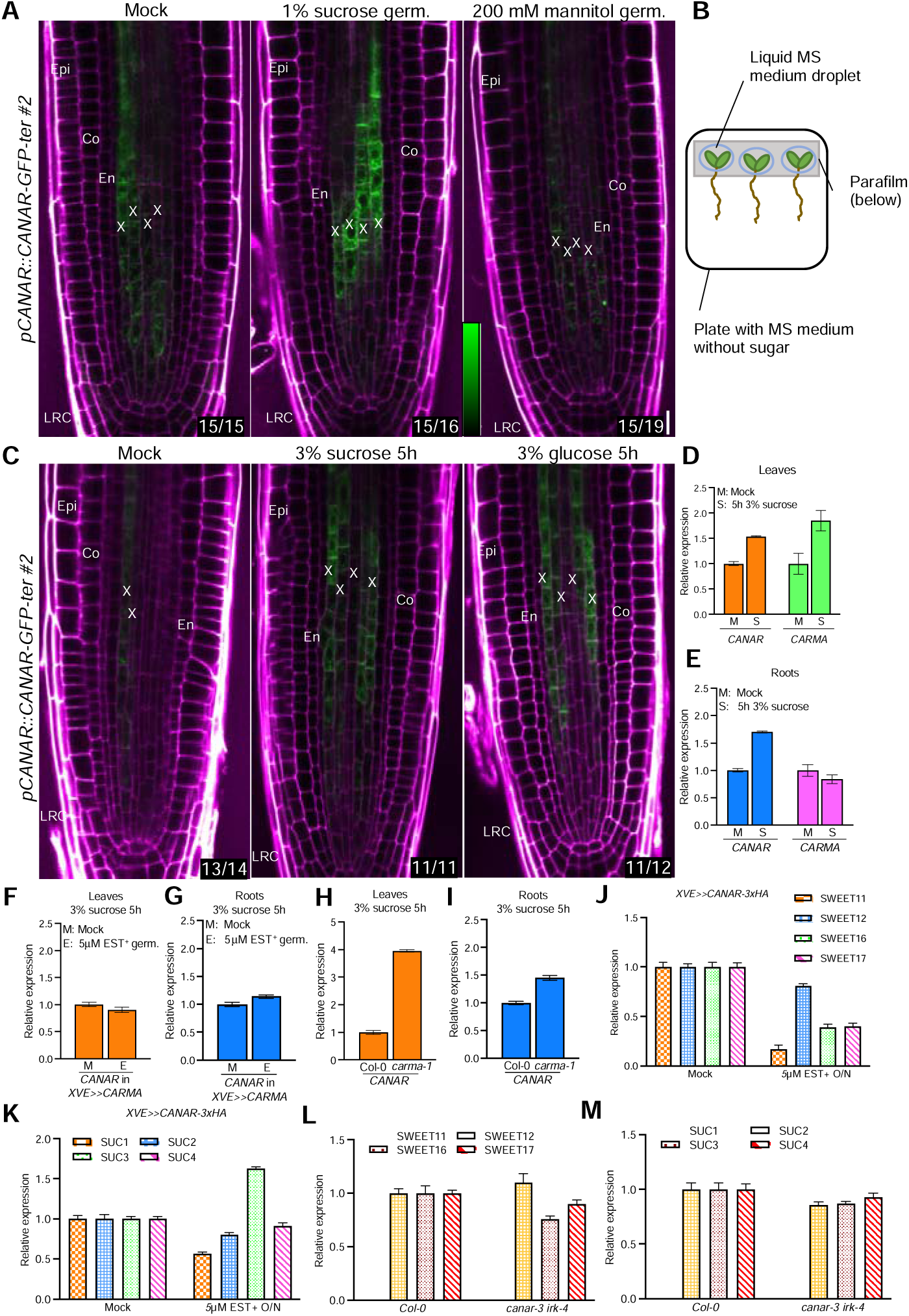
*CARMA* mediates the sugar responsiveness of CANAR. (A) Representative confocal images of primary roots stained with propidium iodide (magenta) expressing *pCANAR::CANAR-GFP-ter* on 0.5x MS medium and medium with sucrose or mannitol. (B) Schematic representation of experiment in (C). (C) Representative confocal images of primary roots stained with propidium iodide (magenta) expressing *pCANAR::CANAR-GFP-ter* 5-hours after application of droplets of liquid 0.5x MS medium containing sucrose or glucose to the shoots. For each treatment, ≥12 roots were analyzed, and the images were acquired using comparable settings. Scale bar, 20 μm. Cell types: Epi-epidermis, Co-cortex, En-endodermis, X-xylem, and LRC-lateral root cap. White numbers at the bottom right corner indicate a frequency of observed expression pattern. (D) and (E) RT-qPCR expression analysis of *CANAR* and *CARMA* after 5-hour sucrose treatment in liquid 0.5x MS media. (F) to (I) RT-qPCR expression analysis of *CANAR* in *XVE>>CARMA* and *carma-1* plants after sucrose treatment in liquid 0.5x MS media. (J) to (M) RT-qPCR expression analysis of *SWEET11/12/16/17* and *SUC1/2/3/4* sucrose transporters in *XVE>>CANAR-3xHA* and *canar-3 irk-4* plants. The graphs show data from three biological replicates. Error bars represent SE.

Further, we tested if CANAR expression in the root could respond to sugars transported from the shoot. Plants were grown on 0.5x MS medium without sucrose for five days, and then the shoots were placed on parafilm to separate them from the media. Then, the shoots were exposed to liquid 0.5x MS medium alone or containing sucrose or glucose (Fig. 4B). After five hours, we observed CANAR upregulation in the root (Fig. 4C).

Increased CANAR accumulation in the root upon exposure to sucrose is, at least partially, explained by increased *CANAR* mRNA in leaves and roots (Fig. 4D, E). This upregulation is impaired when *CARMA* is overexpressed (Fig. 4F, G), and in *carma-1, CANAR* expression upon sucrose treatment is enhanced in leaves (Fig. 4H, I). Indeed, *CARMA* itself is upregulated by sucrose only in leaves (Fig. 4D, E), which could clarify the former observation.

In summary, sucrose upregulates *CANAR* expression, and this effect appears to be mediated by *CARMA*. The upregulation is specific to PM-permeable sugars since using other osmotically active molecules did not mimic the effect.

### CANAR regulates the expression of sugar transporters

The upregulation of CANAR in response to sugars led us to hypothesize that CANAR may regulate sugar distribution. In line with our hypothesis, inducing *CANAR* overexpression in *XVE>>CANAR-3xHA* seedlings by growing them on 0.5x MS media with β-estradiol strongly reduced growth (Fig. S6A, C). This pleiotropic phenotype is reminiscent of various sugar transporter mutants or overexpression lines (*18*). This phenotype was partially rescued by external sucrose application (Fig. S6B, D). Therefore, we examined the expression of sugar transporters in plants overexpressing *CANAR*. SWEETs have been most extensively characterized in *Arabidopsis thaliana,* which contains four SWEET clades: I and II for final distribution of sucrose, glucose, and fructose within sink tissues, III for phloem loading and unloading, and IV for vacuolar sugar storage (*18*). Additionally, Arabidopsis contains nine SUC transporters (SUC1-9) (*19*). We selected *SWEET11*/*12,* which are expressed in leaf phloem parenchyma cells and affect vascular development (*20*), and *SWEET16/17,* which function in root vacuolar storage of glucose and fructose (*21*). For the *SUCs*, we chose *SUC1/2/3/4,* which are expressed in the shoot and root, with SUC2 being the main contributor to shoot-to-root sucrose transport (*22*). We induced *CANAR* expression overnight to allow for sufficient protein translation while avoiding secondary effects from prolonged treatment. All tested *SWEETs,* except *SWEET12,* were strongly downregulated (Fig 4J). *SUC1/2/4* were downregulated as well, while *SUC3* was upregulated (Fig. 4K). In a complementary experiment, we tested *SWEET* and *SUC* expression in the *canar-3 irk-4* double mutant. We found that *SWEET11/16/17* were downregulated, and *SUC1* and *SUC2/3/4* were slightly upregulated and downregulated, respectively (Fig. 4L, M). We did not observe any pronounced effect in the *canar-3* single mutant, which aligns with its reported redundancy with IRK (*10*). Moreover, tissue-specific effects may be concealed due to the inherently low resolution of RT-qPCR with whole seedlings. These results indicate that sugar transporters are downstream of the activity of the CARMA-CANAR module.

## Discussion

Here, we propose that the CARMA-CANAR module acts as a novel osmoregulatory system controlling cell size in the stele. We conclude that the CANAR function is connected to sugar distribution, which influences water retention and, thus, resultant cell size. A link between subcellular sugar distribution and internal cell pressure was proposed previously (*23*), where the SWEETs and aquaporins in *Setaria viridis* guide sucrose and water partitioning between vacuoles, cytosol, and the storage parenchyma apoplast to adjust cell turgor. We propose that CANAR contributes to water distribution by modifying the expression of *SUCs* and *SWEETs*, but the mechanism is unclear. Given that CANAR is a PM-localized pseudokinase and influences the expression of both PM-and vacuolar-localized sugar transporters, a direct interaction is unlikely. Thus, it is more plausible that CANAR controls some aspects of the sugar transporters’ direct regulator/s. Notably, sugar distribution is an intricate system involving various transporters needed to transport sugars between different subcellular compartments and enzymes required to interchangeably convert sugars from one form to another based on *in planta* demand. This complexity is understandable, given the central role of sugars in plant growth and development. For additional perspective, the *SWEET* family in Arabidopsis contains 20 genes, whereas animal genomes have only one (*5*). Moreover, the exact molecular function of sugar transporters in phloem loading/unloading is not entirely clear. We, therefore, cannot distinguish the direct effect of *CARMA-CANAR* manipulation on sugar distribution from a compensatory response of the plant.

We reveal the long non-coding RNA *CARMA* modulates the sucrose-mediated upregulation of *CANAR* in the protophloem. Fine-tuned levels of protophloem-expressed *CANAR* are necessary for the adaptation of cell size to external osmolality, which ensures the maintenance of optimal stele area. It is still unclear what CANAR’s role in the xylem is. Water exchange between the xylem and phloem is necessary to load and transport phloem content (*2*). Xylem-expressed CANAR could potentially help to facilitate this exchange.

Our hypothesis about the osmoregulatory function of the CARMA-CANAR module may explain the extra endodermal divisions in the *irk-4* and *canar-3 irk-4* mutants and their absence in *canar-3* (*9*, *10*). Larger cells in the stele generate elevated mechanical pressure on the endodermis, the pressure-buffering tissue of the root (*8*). Both *canar-3* and *irk-4* plants have an enlarged stele area, although the increase is greater in *irk-4*. This suggests there is a certain pressure threshold after extra divisions in the endodermis are induced as a coping mechanism to dissipate the built-up mechanical pressure in the stele. This hypothesis is corroborated by the *canar-3 irk-4* double mutant, in which the stele area was more enlarged than in the single mutants, resulting in a higher incidence of extra endodermal divisions compared to *irk-4* (*10*). In line with our observations, a cellulose-deficient *korrigan-1* mutant displayed root thickness twice that of the wild type (*24*). The enlargement resulted from all examined cell types (epidermis, cortex, endodermis, and pericycle), with the greatest contribution from cortex cells. Swollen cortex cells generated mechanical pressure towards the outer epidermal cells and cells of inner tissues. Still, mechanical stress, as evidenced by elevated jasmonate signaling, was observed only in endodermal and pericycle cells. The authors reasoned that epidermal cells dissipated the excessive pressure by expanding outward into the rhizosphere, and no extra cell divisions were induced in the endodermis.

The observations that *IRK* mutant (*9*) and *CARMA* and *CANAR* (*11*) mutant/overexpressing lines exhibit defects in leaf vascular patterning suggest that stele area and leaf vein patterning (via auxin canalization) are developmentally co-dependent. It is possible that an appropriate stele area is required for undisturbed vascular patterning or that sugars are vital signaling molecules instructing auxin canalization and, thus, vasculature establishment. However, we cannot uncouple these two phenomena as the vasculature in cotyledons is already established in the embryo. Both scenarios are plausible as mechanical signals (laser ablation) in the shoot meristem induce reorientation of PIN1 auxin exporter (*25*), and leaf vasculature still forms, although imperfectly when auxin directional transport is not functional (*26*). Perhaps the residual vein-patterning activity could be attributed to the sugar transport? Alternatively, SWEET transporters might transport auxin, as it was recently reported that *Arabidopsis* SWEET13/14 proteins can transport multiple forms of gibberellins (*27*). This broad substrate specificity is also displayed by ABCB transporters that contribute to directional auxin transport (*28*).

Besides the energy value of sugars, they also serve as signaling molecules. An extensive sugar-auxin signaling interaction network was recently described (*29*). For instance, high glucose levels increased PIN2-GFP accumulation at the PM, promoting basipetal auxin transport in *Arabidopsis* (*30*) while compromising PIN1-GFP expression, reducing auxin concentration in the root tip (*31*). Given the interaction of CANAR with PIN1 (*11*), the CARMA-CANAR module could be involved in the intricate interplay between sugar-auxin.

In the root, the endodermis, xylem, and protophloem are interconnected in pressure sensing. Furthermore, the mature endodermis is an essential selective barrier restricting the free diffusion of solutes into vascular tissues (*32*). Further work is needed to elucidate how IRK/CANAR function to modulate stele area and endodermal cell division and how they are mechanistically linked to sugar distribution. Manipulation of stele width and sugar distribution would have promising benefits for agriculture, for example, removing sugars from the extracellular space (apoplast) reinforces the defense against microbial infection (*4*), and sugar content also changes plant susceptibility to drought, cold, and heat stress (*5*). However, progress is hindered by a lack of known molecular regulators, and our work may provide key entry points into these important questions.

## Material and Methods

### Plant Materials and Growth Conditions

All *Arabidopsis thaliana* lines were in Columbia-0 (Col-0) background. The T-DNA insertional mutant of *carma-1* (SAIL_704_A04) was obtained from NASC and genotyped with the primers listed in Supplemental Table 3. The *canar-3* (*pxc2-3*, SM_3_31635), *canar-3/pxc2-3 irk-4*, and *XVE>>CANAR* were described previously (*10*). Seeds were sterilized with 70% ethanol for 5 min and then with 100% ethanol for another 5 min. Seeds were plated on 1% plant agar pH 5.9 (Duchefa) supplemented with 0.5x Murashige and Skoog (0.5x MS) media basal salts (Duchefa) unless otherwise indicated. 5-days old seedlings were used for imaging (counting 5 days after placement in the Phytochamber) When testing the effect of sugar, plants were grown from germination on 0.5x MS with 1% sucrose or incubated for 5h in liquid 0.5x MS with 3% sucrose or glucose. Transgenic lines with the ®-estradiol inducible promoter (*XVE*) were grown on 5 ⎧M ®-estradiol from germination unless otherwise indicated. Plates were sealed with 3M micropore tape. Seeds were stratified on plates at 4°C for 1-2 days before being placed in a Phytochamber (16h light/8h dark cycle at a constant temperature of 21°C, light intensity ∼ 700-foot candle).

### Cloning and Plant Transformation

Transcriptional reporter for *CANAR* (AT5G01890) was constructed by LR recombination of 4.7 kb promoter in pENTR5‘-TOPO (*10*) with NLS-GFP-GUS and 285 bp of *CANAR* 3’UTR region (*ter*) in pENTR2B (generated via Gibson assembly-NEBuilder Hifi DNA assembly Master Mix) into pK7m24GW-FAST destination vector. The deletion of 157 bp of *CARMA* (until annotated 5’ UTR of *CANAR*) was performed by amplifying truncated *pCANAR* in pENTR5‘-TOPO with primers containing a SalI restriction site. The amplicon was cut with SalI for 30 min (FastDigest; Thermo), cleaned, and ligated overnight at 16°C (T4 DNA ligase; NEB). The same approach was used for the second deletion (353 bp) of the *CARMA* locus. All three versions: *pCANAR::NLS-GFP-GUS-ter*, *pCANAR_CARMA*Δ*::NLS-GFP-GUS-ter* and *pCANAR_CARMA*ΔΔ*::NLS-GFP-GUS-ter* were transformed into *carma-1* (SAIL_704_A04). Transcriptional reporters for *CARMA* (AT5G00810) were constructed by inserting 1300 bp *CARMA* promoter into pDONRP4-P1R via BP reaction and inserting 4975 bp *CARMA* promoter into pENTR5’TOPO via Gibson assembly. pDONRP4-P1R was recombined into pMK7S*NFm14GW, and pENTR5’TOPO with NLS-GFP-GUS in pENTR2B (NLS-GFP-GUS fragment was amplified from pMK7S*NFm14GW and inserted in pENTR2B via SalI restriction and subsequent ligation) into pH7m24GW destination vector via LR reaction. Translation reporters were constructed using Invitrogen Multisite Gateway technology. *pCANAR* (in pENTR 5’ TOPO), *pCVP2-XVE* (in pDONRP4-P1R) were recombined with *CANAR* (genomic fragment without stop codon in pENTR-D-TOPO) (*10*) and with *GFP-ter* (GFP flanked by pkpapkpa linker at N-terminus and *CANAR* 285 bp 3’ UTR region at C-terminus in pDONRP2r-P3) via LR reaction. For a generation of *XVE>>CARMA*, the genomic fragment of *CARMA* (AT5G00810) was amplified from Col-0 genomic DNA and recombined into the pDONRP221 entry vector via BP reaction. This was then recombined into the pMDC7 destination vector via LR reaction. All primers used are listed in Supplemental Table 3.

### Plant transformation

Transgenic *Arabidopsis thaliana* plants were generated by the floral dip method using *Agrobacterium tumefaciens* (strain GV3101).

### RNA extraction, cDNA synthesis, and quantitative RT-PCR analysis

Total RNA was isolated from seedlings for gene expression analysis in mutants and overexpressing lines or from roots for RNA sequencing using Spectrum Plant Total R.N.A. Kit (Sigma). RNA was treated with TURBO DNAse (Thermo) to avoid genomic DNA contamination. Three independent biological replicates were done per sample. For cDNA synthesis (RevertAid First Strand cDNA Synthesis kit, Thermo), 2 μg of total RNA was used with Random Hexamer Primers mix (for RT-PCR of *CARMA* in Fig.1D) or with Oligo(dT) for the rest of the RT-qPCRs. The generated cDNA was analyzed on the StepOnePlus Real-Time PCR system (Life Technologies) with gb SG PCR Master Mix (Generi Biotech) according to the manufacturer’s instructions. The relative expression was normalized to *SERINE/THREONINE PROTEIN PHOSPHATASE, PP2A (AT1G69960)*. Three technical replicates were performed.

### Confocal microscopy

Five days-old roots were stained with propidium iodide (PI) (10 µg/mL) and visualized via laser scanning confocal microscopy using a Zeiss LSM900 with a 40x water immersion objective. Fluorescent signals were visualized as PI (excitation 536 nm, emission 585-660 nm) and eGFP (excitation 488 nm, emission 492-530 nm). For stele area analysis, Z-stacks of approximately 100 μm were taken. ImageJ software was used for image postprocessing and quantification of stele area.

### Histological analyses

β-glucuronidase (GUS) staining was performed as described in (*33*). The staining reaction was stopped with 70% ethanol and left for two days to remove chlorophyll. Seedlings were mounted in chloral hydrate and examined using a stereomicroscope (Olympus). ClearSee tissue clearing (*34*) was performed to count the cells in the transverse optical sections. The seedlings were fixed in 4% PFA in PBS (1h in vacuum), washed with PBS, and placed into ClearSee solution (25% urea, 15% sodium deoxylate, and 10% xylitol) for at least 3 days. Then, the seedlings were transferred into 0.1% Calcofluor White in ClearSee solution for 60 min, followed by a wash with ClearSee solution for 30 min; then mounted on slides with ClearSee. Two-sided tape was used on slides to prevent tissue disruption.

### Stele area and vascular cell number quantification

Z-stacks of ∼100 μm (1μm thick slices) capturing the root meristematic zone were acquired. Bleach correction plugin in ImageJ was applied to all images to compensate for decreasing PI signal in the deeper part of the root. The stele area and the number of vascular cells were assessed in the transverse sections located ∼100 μm above QC using ImageJ.

### Quantification of pCANAR expression in protophloem

Z-stacks of approximately 100 μm capturing the root meristematic zone was acquired. Multiple transverse sections with nuclear GFP fluorescence in xylem and protophloem in the same plane were taken for each Z-stack. The fluorescent signal in the protophloem was normalized to the xylem signal in each transverse section, and the average value of all sections from one root was calculated and plotted into a graph.

### Software

Postprocessing of confocal images was done in ImageJ (https://imagej.nih.gov/ij/). Figures were generated in Adobe Illustrator. Graphs and statistics were completed in GraphPad Prism9.

### 5 ‘and 3 ‘RACE experiments

The 5’RACE-seq library was generated from five-day-old roots with template-switching RT following the protocol outlined in (Montez et al, 2023). Shortly, 500 ng of total RNA, post DNAse treatment, served as the template for cDNA generation using SuperScript II. The resulting cDNA was purified using AMPure XP magnetic beads (Beckman Coulter) and amplified in series of three PCR reactions with specific primers (1^st^ PCR: only TSO_n1, 2^nd^ PCR: TSO_n2 and CARMA_5RACE, 3^rd^ PCR: Illumina indexing primers) and Phusion polymerase. Following quality checks, the final PCR product was sequenced using Illumina MiSeq.

The 3’RACE-seq was completed based on the procedure described by Warkocki et al (2018) with ligation of the pre-adenylated adaptor to the 3’end of the RNA using truncated T4 RNA Ligase 2. RNA ligated with RA3_15N adaptor (containing UMI) was cleaned on AMPure XP magnetic beads and subjected to RT reaction with SuperScrit III. After three rounds of PCR with specific primers (1^st^PCR: CARMA_3RACE and RTPXT, 2^nd^PCR: mXTf and mXTr, 3^rd^ PCR: Illumina indexing primers) and cleaning each PCR reaction on AMPure beads, prepared libraries were sequenced using Illumina MiSeq.

Sequence reads were trimmed to remove adapter sequences using cutadapt (v1.18; Martin, 2011). STAR (v2.7.8a; Dobin et al, 2013) was utilized to align the reads to the reference genome, followed by UMI-based filtering using UMI-tools (v1.1.0; Smith et al, 2017). The position of the reads ends nucleotide was extracted using bedtools (v2.30.0; Quinlan & Hall, 2010).

## Acknowledgments

We acknowledge the EMBO long-term fellowship (ALTF 217-2021) and Junior grant UPOL (JG_2024_003) for supporting JH. The work of RMIKR and JMVN is supported by NSF CAREER award #1751385.

## Author contributions

Conceptualization: JH; Resources: ON; Writing, editing and interpretation of data: JH, SS, SS, AP, JMVN, DZ, CH, RMIKR; Methodology: JH, DZ, SS, TT, RMIKR; RACE experiments: SS, SS; Bioinformatics: MK, Imaging: JH, RMIKR; Cloning: JH, DZ, TT; Generation of transgenic lines: JH, TT.

## Declarations

The authors declare no conflict of interest.

---

**Supplemental Fig. 1 Characterization of the *CARMA* transcript**

(A) and (B) The full-length transcript of *CARMA* based on 5’ and 3’ RACE results. Representative confocal images of a primary root stained with propidium iodide (grey) of roots showing expression of (C) *pCARMA(5kb)::NLS-GFP-GUS* and (D) *pCARMA(1.3kb)::NLS-GFP-GUS* (depicted above images). (E) *pCARMA* activity visualized by β-glucuronidase (GUS) staining in a root expressing *pCARMA(1.3kb)::NLS-GFP-GUS*. A minimum of 10 roots were examined for each reporter. Scale bars, 20 μm. Cell types: Epi-epidermis, Co-cortex, En-endodermis, PPh-developing protophloem sieve elements, MPh-metaphloem precursors, X-xylem, LRC-lateral root cap.

**Supplemental Fig. 2 Enlarged stele area phenotype upon *CARMA* overexpression is due to larger cells.**

(A) The position of T-DNA insertion in the *carma-1* mutant. (B, D, E) Relative expression by RT-qPCR of *CARMA* in Col-0, *35S::CARMA, XVE>>CARMA* and *carma-1.* (C) Relative expression by RT-qPCR of *CANAR* in *carma-1*. The graphs represent three biological replicates. Error bars represent SE. (F) Distance between endodermis and lateral root cap in *XVE>>CARMA* line as visualized in (H) by the orange bidirectional arrow. The experiment was carried out three times (each with 10 roots per sample per genotype), data shown are from a single biological replicate. (G) The number of cells in stele in *XVE>>CARMA* with and without β-estradiol induction from the time of germination with 15-20 roots analyzed per line per condition. (H) Representative transverse optical sections taken ∼100 µm from QC (quiescent center), where stele area was quantified for (G). (J) Box plot showing root meristem lengths from (I). Whiskers indicate the max/min, box shows interquartile range, and the median is shown with a black line. These analyses were performed three times with ≥18 roots per genotype per condition. Graphs show the data from 1 biological replicate. (I) Representative images of the median longitudinal sections of the root meristem of two independent *XVE>>CARMA* transgenic lines with and without β-estradiol from the time of germination on 0.2x MS medium.

**Supplemental Fig. 3 Reduced stele area phenotype in *carma-1* is due to smaller cells.**

(A) Number of cells within the stele in *carma-1* mutants compared to Col-0 (n = X). (B) Representative transverse sections taken approximately 100 µm from QC (quiescent center) where stele cells were counted for (A) with 15-20 roots analyzed. (C) Measurement of the distance between the endodermis and LRC in Col-0 and *carma-1* on 1x MS media. (D) Representative images of the median longitudinal sections of the Col-0 and *carma-1* root meristems. (E) Measurement of meristem length in Col-0 and carma-1 on 1x MS medium. These analyses were carried out three times with ≥15 roots per genotype. Graphs show the data from 1 biological replicate. Scale bars, 20 μm. Cell types: Epi-epidermis, Co-cortex, En-endodermis, Per-pericycle, X-xylem.

**Supplemental Fig. 4 *CARMA* regulates the protophloem-specific expression of *CANAR***

(A) Representative confocal images of primary roots stained with propidium iodide (white) of a second independent transgenic line of each *pCANAR::NLS-GFP-GUS-ter, pCANAR_CARMA*Δ*::NLS-GFP-GUS-ter,* and *pCANAR_CARMA*ΔΔ*::NLS-GFP-GUS-ter* in *carma-1* (schematics of each reporter above the images). Both partial and complete deletion of *CARMA* show increased *pCANAR* activity in the PPh (highlighted with green text and arrowhead). Numbers represent the position of a transverse optical section taken from Z-stacks.

(B) Box plot shows the quantification of fluorescent signal from (A), where signal from the PPh is normalized to that from the X (see the Material and Method section for details). Whiskers indicate the max/min with boxes showing interquartile range, and a black line shows the median. Colored symbols indicate measurements from individual roots. These experiments were done three times (8-10 roots for each genotype per experiment); one representative biological replicate is shown. A one-way ANOVA test compared marked datasets (***P<0.001). Scale bar, 20 μm. Cell types: Epi-epidermis, Co-cortex, En-endodermis, PPh-developing protophloem sieve elements, X-xylem, LRC-lateral root cap.

**Supplemental Fig. 5 CANAR is specifically upregulated by PM-permeable sugars**

(A) Representative confocal images of primary roots stained with propidium iodide (magenta) of a second, independent *pCANAR::CANAR-GFP-ter* line on 0.5x MS medium and medium with sucrose or mannitol. (B) Representative confocal images of *pCANAR::CANAR-GFP-ter* after 5-hour treatment with liquid 0.5x MS alone or with sucrose or glucose. Scale bar, 20μm. (C) Representative confocal images of roots expressing *pCANAR::CANAR-GFP-ter* after 5-hour treatment with liquid 0.2x MS, 0.5x MS, 1x MS, and 0.5x MS with NaCl. White numbers in the bottom right corner indicate a frequency of observed expression pattern. (D) RT-qPCR expression analysis of *SWEET11/12/16/17* and *SUC1/2/3/4* sucrose transporters in the *canar-3* mutant. The graphs show data from three biological replicates, and error bars represent SE.

**Supplemental Fig. 6 *CANAR* overexpression leads to reduced seedling growth that can be rescued by exogenous sucrose**

(A, B) Representative images of six-day-old seedlings expressing *XVE>>CANAR-3xHA* that are mock-or β-estradiol-treated grown on (A) 0.5x MS media and (B) 0.5x MS media with 1% sucrose. (C, D) Box plots show root length quantification from (A) and (B). Whiskers indicate the max/min, boxes indicate interquartile range, and the median is shown with a black line. Colored symbols show measurements from individual roots. These experiments were carried out three times; one representative biological replicate is shown (n is approximately 80 roots per replicate per genotype). A one-way ANOVA test compared marked datasets (****P<0.0001). Scale bar, 10 mm.

